# Isolation of bat sarbecoviruses of SARS-CoV-2 clade, Japan

**DOI:** 10.1101/2022.05.16.492045

**Authors:** Shin Murakami, Tomoya Kitamura, Hiromichi Matsugo, Haruhiko Kamiki, Ken Oyabu, Wataru Sekine, Akiko Takenaka-Uema, Yuko Sakai-Tagawa, Yoshihiro Kawaoka, Taisuke Horimoto

## Abstract

Betacoronaviruses have caused 3 outbreaks in the past 2 decades. SARS-CoV-2, in particular, has caused a serious pandemic. As the betacoronaviruses are considered to originate from bats, surveillance of bat betacoronaviruses is crucial for understanding the mechanism of cross-species transition and potential for future outbreaks. We previously detected and characterized a SARS-CoV-2-related sarbecovirus, Rc-o319, from *Rhinolophus cornutus* in Japan. Here, we detected several bat sarbecoviruses of the SARS-CoV-2 clade from *R. cornutus* in multiple locations in Japan, and successfully isolated them using Vero/TMPRSS2 cells stably expressing *R. cornutus* ACE2 (Vero-RcACE2). The coding sequences of S1 region varied among isolates, whereas other genetic regions were highly conserved. Isolates were efficiently grown in Vero-RcACE2 cells, but did not replicate in Vero/TMPRSS2 cells stably expressing human ACE2, suggesting a narrow host range. Further long-term epidemiological studies of sarbecoviruses in wildlife are expected to facilitate the assessment of the risk of their spillover potential.

---

Severe acute respiratory syndrome coronavirus (SARS-CoV), Middle East respiratory syndrome coronavirus (MERS-CoV), and SARS-CoV-2 have caused 3 outbreaks in human populations in the past 2 decades. In particular, SARS-CoV-2 has caused a worldwide serious pandemic with devastating damage to human health, many social activities, and the economy. All these betacoronaviruses are considered to be of bat origin. Hence, surveillance of bat betacoronaviruses is crucial for understanding and assessing the spillover potential of betacoronaviruses in humans in the future.

SARS-CoV and SARS-CoV-2 belong to the *Betacoronavirus* genus and *Sarbecovirus* subgenus. Bats belonging to the genus *Rhinolophus* are considered natural reservoirs of sarbecoviruses, as most have been detected in this bat species in Asian countries such as China (1-9), Japan (10), South Korea (11, 12), Laos (13), Thailand (14), and Cambodia (15), as well as in European and African countries such as Bulgaria (16), England (17), and Kenya (18). Bat sarbecoviruses closely related to SARS-CoV-2 have also been found in the Yunnan province, China, and Southeast Asian countries (1, 3, 7, 8, 13-15) with some possessing the ability to bind to the human angiotensin-converting enzyme 2 (ACE2) as an entry receptor, posing a potential for human infection (3, 8, 13).

We previously identified a Japanese bat sarbecovirus, Rc-o319, detected from *Rhinolophus cornutus* bats in the Iwate prefecture, which was shown to phylogenetically belong to the SARS-CoV-2 lineage. Vesicular stomatitis virus (VSV)-based pseudotyped virus possessing the Rc-o319 spike (S) protein was able to infect cells expressing *R. cornutus* ACE2 (RcACE2), but not those expressing human ACE2 (hACE2), suggesting that the Rc-o319 virus uses RcACE2 as its receptor (10). Sarbecoviruses detected in China and other Asian countries were demonstrated to vary genetically. However, the distribution and genetic variation of bat sarbecoviruses in Japan have not yet been determined.

Despite the surveillance-based genetic detection of numerous bat sarbecoviruses, infectious viruses have been limitedly isolated to date. Infectious viruses are useful and required for understanding the biological characteristics. Here, we report the detection and isolation of bat sarbecoviruses from several locations in Japan and their genetic and biological characterization.

## Materials and methods

### Cells and virus

Vero/TMPRSS2 cells (19) were kindly provided by Dr. Makoto Takeda, National Institute of Infectious Diseases, Japan, and maintained in Dulbecco’s modified Eagle’s medium (DMEM, Nacalai Tesque, Kyoto, Japan) supplemented with 10 % fetal bovine serum (FBS), 100 units/mL of penicillin, and 100 μg/mL of streptomycin. Cells were cultured in 5 % CO2 at 37 °C. SARS-CoV-2 (B.1.1.7, alpha variant, UT-HP127-1Nf/Human/2021/Tokyo) was propagated in Vero/TMPRSS2 cells, and aliquots were stored at −80 °C.

### Sample collection

We collected 88 fresh fecal samples, 58 from *Rhinolophus cornutus* and 30 from *Rhinolophus ferrumequinum* bats living in caves, abandoned mines, or abandoned tunnels in Niigata, Chiba, and Shizuoka prefectures in Japan (Table 1). When bats were densely packed during daytime roost, plastic sheets were placed under the roost for 1-2 h. Fresh feces that dropped onto the sheets were collected. When bats were sporadically placed during daytime roost, we captured bats with permission from the prefectural local governments (No. 962 for Chiba and No. 311 for Shizuoka) and kept each bat in a separate nonwoven fabric bag. Feces excreted by the bat in the bag were collected and bats were released. Fecal samples were transferred into tubes containing phosphate-buffered saline (PBS) supplemented with 200 U/mL penicillin, 200 μg/mL streptomycin, and 0.25 μg/mL amphotericin B, and immediately frozen in dry ice.

**Table 1.** Detection of sarbecoviruses in Japan using RT-PCR.

| Location | Bat species | Number of samples | Number of positive RT-PCR samples* |
| --- | --- | --- | --- |
| Niigata | *R. cornutus* | 26 | 2 |
|  | *R. ferrumequinum* | 1 | 0 |
| Chiba | *R. cornutus* | 11 | 1 |
|  | *R. ferrumequinum* | 16 | 0 |
| Shizuoka | *R. cornutus* | 21 | 2 |
|  | *R. ferrumequinum* | 13 | 0 |

\*RT-PCR was performed using sarbecovirus consensus primers targeting the E gene.

### Reverse transcription-PCR

RNA was extracted from the fecal samples using the RNeasy PowerMicrobiome Kit (Qiagen KK, Tokyo, Japan) and the partial E gene of sarbecovirus was detected in RNA samples by real-time reverse transcription-PCR (rRT-PCR) using the RNA-direct SYBR Green Realtime PCR Master Mix (Toyobo, Osaka, Japan) and a pair of primers (5′-TCGGAAGAGACAGGTACGTT-3′ and 5′-TCGAAGCGCAGTAAGGATGG-3′) that were designed to target a highly conserved region of the sarbecovirus E gene.

### Establishment of ACE2-stably expressing cells

We constructed a plasmid, pCAGGS-blast, by inserting the XhoI-EcoRV fragment of pMXs-IRES-Bsd (20), which contains the encephalomyocarditis virus internal ribosomal entry site and blasticidin-resistant gene, into the XhoI and StuI sites of the pCAGGS-MCS vector. Open reading frame (ORF) sequences of RcACE2 or hACE2 were PCR-amplified from pCAGGS-RcACE2- or pCAGGS-hACE2-expressing plasmids (10) and cloned into EcoRI- and XhoI-digested pCAGGS-blast plasmids using NEBuilder (New England Biolabs, Ipswich, MA, USA). Vero/TMPRSS2 cells were transfected with pCAGGS-blast-RcACE2 or pCAGGS-blast-hACE2 plasmids using the PEI MAX transfection reagent (Polysciences, Warrington, PA, USA). Transfected cells were treated with 10 μg/mL blasticidin S (Kaken Pharmaceutical, Tokyo, Japan) 1 d posttransfection, and blasticidin S-resistant cells were selected and cloned. Highly susceptible cell clones for the pseudotyped viral infection were selected by screening using GFP-expressing VSV-pseudotyped virus possessing the S protein of Rc-o319 or SARS-CoV-2 (10), generating RcACE2- or hACE2-stably expressing Vero/TMPRSS2 cells (namely Vero-RcACE2 or Vero-hACE2, respectively).

### Establishment of ACE2-knockout cells

We generated ACE2-knockout Vero/TMPRSS2 cells (Vero-ACE2KO) by knocking out the corresponding genes using the CRISPR/Cas9 system. The target sequence for the ACE2 gene (5′-TGCTGCTCAGTCCACCATTG-3′) was designed using CRISPR direct (https://crispr.dbcls.jp) and cloned into plentiCRISPR plasmids (21) (Addgene plasmid #52961, a gift from Dr. Feng Zhang) using NEBuilder (NEB). Vero/TMPRSS2 cells were transfected with an ACE2-targeting plasmid using PEI MAX (Polysciences). At 24 h posttransfection, the cell supernatant was replaced with medium containing 10 μg/mL puromycin. Drug-resistant clones were randomly selected and their genomic DNA was sequenced. Cells possessing insertions or deletions (in/dels) in the targeted gene were chosen for further analysis.

### Isolation of bat sarbecoviruses

Fecal samples positive for the partial E gene of sarbecovirus were homogenized in TissueLyser II (Qiagen) using 0.1 mm glass beads (Tomy Seiko, Tokyo, Japan) in PBS containing 200 U/mL penicillin, 200 μg/mL streptomycin, and 0.25 μg/mL amphotericin B. The supernatants were collected after centrifugation at 5000 × g for 5 min at 4 °C and diluted 100-fold in cell maintenance medium (DMEM supplemented with 1 % FBS, 200 U/mL penicillin, 200 μg/mL streptomycin, and 0.25 μg/mL amphotericin B). Diluents were inoculated into 6-well plates containing Vero-RcACE2 cells, and plates were incubated for 60 min at 37 °C after removing the inoculum. Wells were then washed once with cell maintenance medium, another 2 mL of cell maintenance medium was added to each well, and incubated at 37 °C. Supernatants from cells that exhibited cytopathic effects (CPE) were collected at 3–4 d postinoculation and were passed through a 0.22 μm filter. The successful isolation of viruses was confirmed by rRT-PCR, and isolates were propagated in Vero -RcACE2 cells, with aliquots being stored at −80 °C.

### Next-generation sequencing

A cDNA library was prepared from RNA extracted from bat sarbecoviral isolates using the TruSeq Stranded Total RNA LT Sample Prep Kit Gold (Illumina, San Diego, CA, USA) for Rc-mk2 and Rc-kw8 strains or the MGIEasy RNA Directional Library Prep kit (MGI, Shenzhen, China) for the Rc-os20 strain. Libraries of Rc-mk2 and Rc-kw8 strains were sequenced using a Novaseq 6000 sequencer (Illumina), whereas those of Rc-os20 were sequenced using a DNBSEQ-G400RS (MGI) sequencer. Read sequences were mapped to the Rc-o319 genome sequence (GenBank accession No. LC556375) and sarbecoviral sequences were determined using the CLC genomic workbench version 8.0.1 (Qiagen, https://www.qiagen.com) software. The sequences of Rc-os20, Rc-mk2, and Rc-kw8 have been deposited in GenBank (accession Nos. LC663958, LC663959, and LC663793, respectively).

### Phylogenetic analysis

The nucleotide sequences of sarbecoviruses were aligned using ClustalW version 2.1 (Clustal, https://www.clustal.org). Phylogenetic trees were then constructed by performing a maximum-likelihood analysis using MEGA version X (22) in combination with 500 bootstrap replicates.

### Evaluation of viral growth in cells

Cells were inoculated with viruses at a multiplicity of infection (MOI) of 0.01 and allowed for 1 h for viral adsorption. After removing the inocula, cells were incubated in cell maintenance medium, and the supernatants were collected at 12 h intervals. Viral titers were measured using a plaque assay, in which cells inoculated with diluted viruses were overlaid and incubated with DMEM containing 1 % agarose and 1 % FCS for 2 d, followed by staining with crystal violet before counting plaques.

## Results

### Detection of bat sarbecoviruses in Japan

We collected fecal samples from bats belonging to the *Rhinolophus cornutus* and *Rhinolophus ferrumequinum* species in Niigata, Chiba, and Shizuoka prefectures (Figure 1). Using real-time RT-PCR, we successfully detected the E gene sequence of sarbecovirus in 1 or 2 *R. cornutus* samples in each prefecture (Table 1). In contrast, all *R. ferremuquinum* samples were negative. These data suggested that bat sarbecoviruses are distributed among *R. cornutus* at various locations in Japan.

**Figure 1.**
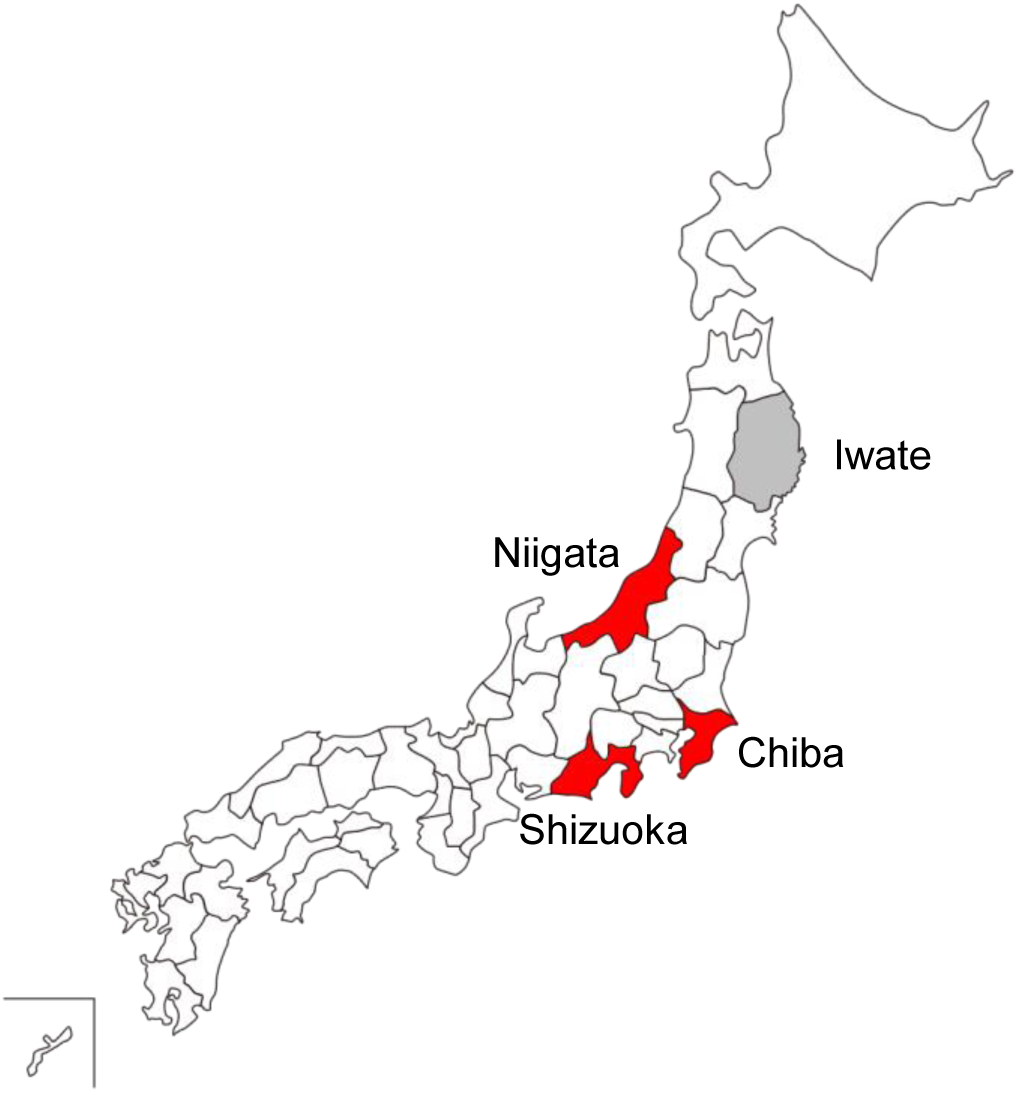
Sampling locations in Japan. Prefectures, where bats were captured, are indicated in red. Prefectures, where bat sarbecoviruses were detected in our previous study, are indicated in grey.

### Isolation of bat sarbecoviruses from *R. cornutus*

In our previous study, we showed that a VSV-based pseudotyped virus possessing the S protein of the Rc-o319 sarbecovirus from *R. cornutus* only infected RcACE2-expressing cells, but not hACE2- or other *Rhinolophus* ACE2-expressing cells (10), suggesting a narrow host range of Rc-o139. Therefore, to isolate bat sarbecoviruses, we established RcACE2-stably expressing cells (Vero-RcACE2) based on Vero/TMPRSS2 cells (19), which are useful for the isolation of SARS-CoV-2 (23). We inoculated Vero-RcACE2 cells with RT-PCR-positive fecal samples from each prefecture and observed any extensive cytopathic effect (CPE) with syncytium formation at 1–2 d postinoculation (Figure 2). Using real-time RT-PCR, we also detected the E gene sequence of sarbecovirus in the supernatants at 3–4 d postinoculation. Following filtration, we inoculated the supernatants into fresh Vero-RcACE2 cells, and observed for CPE, confirming the successful isolation of bat sarbecoviruses. We designated the Niigata, Chiba, and Shizuoka isolates, Rc-os20, Rc-mk2, and Rc-kw8, respectively. Using the same approach, we further isolated the infectious Rc-o319 isolate.

**Figure 2.**
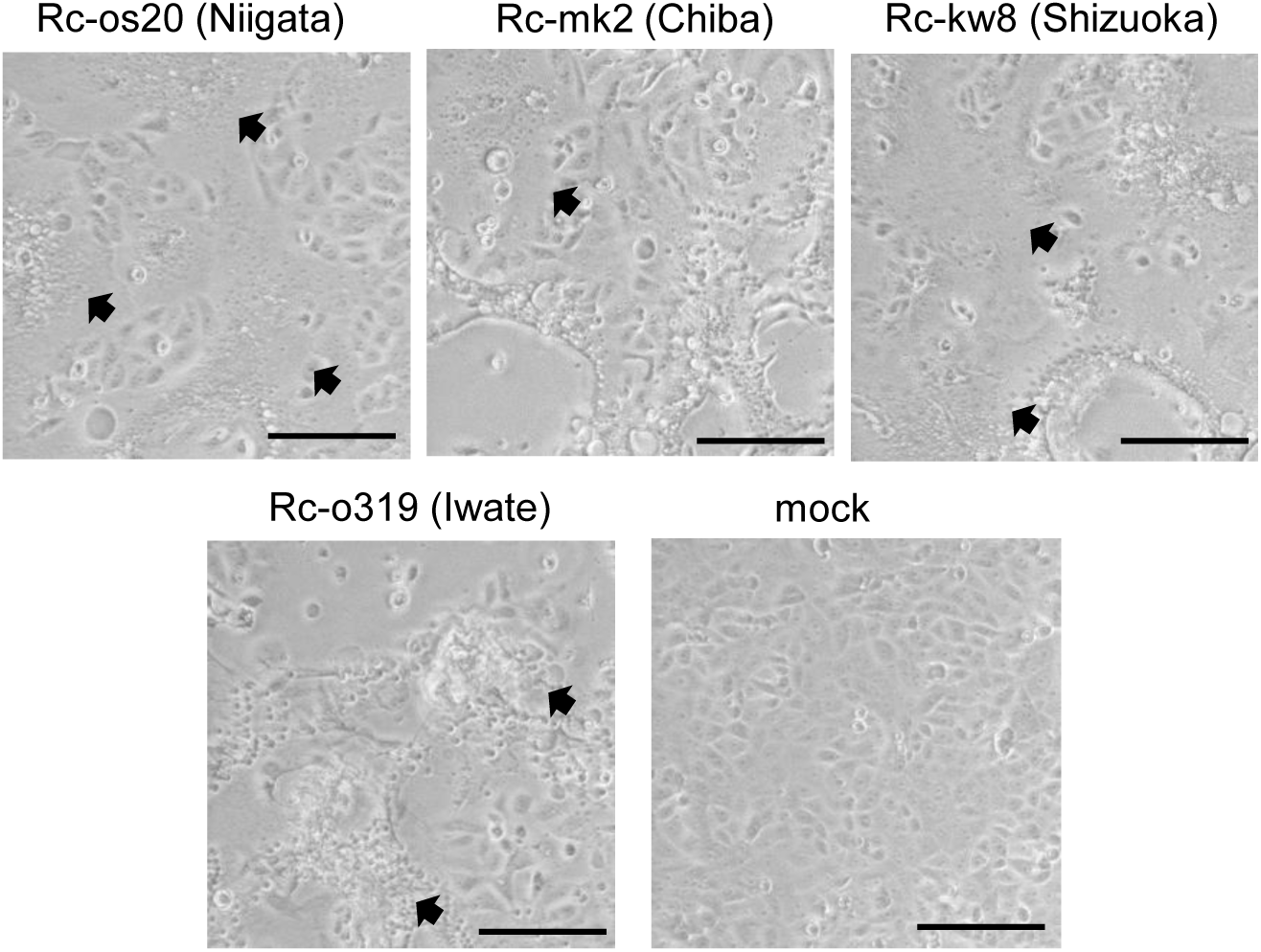
Images of bat sarbecovirus-inoculated cells. Vero-RcACE2 cells were inoculated with fecal samples from *Rhinolophus cornutus* from several prefectures of Japan. After 1–2 d of inoculation, cytopathic effects with extensive syncytium formation (arrow heads) were observed. Scale bar, 200 μm.

### Genetic characterization of Japanese isolates

We determined the full genome sequence of all isolates using next-generation sequencing. We detected that the sequence of Rc-o319 was identical to the previously deposited sequence, which was determined using the Sanger method. Next, we mapped the sequence reads of Rc-os20, Rc-mk2, and Rc-kw8 onto the Rc-o319 genome sequence, and determined their full genomic sequences, which we deposited in GenBank (accession Nos. LC663958, LC663959, and LC663793). We found that sequence homologies were high (ranging 94.8–96.8 %) among all Japanese isolates (Table 2); however, Rc-mk2 and Rc-os20 lacked the entire ORF8 coding region. We also performed similarity plot analysis of entire genome sequence using each isolate as a query, which indicated that the similarities among isolates were high throughout the entire genome sequence, except for coding regions of the N-terminal domain (NTD) and receptor binding domain (RBD) of the S gene, although NTDs of Rc-o319 and Rc-kw8 were conserved (Figure 3A). No clear recombination among the isolates were observed as analyzed by RDP5 software (24). Phylogenetic analysis showed that the Japanese isolates formed a single genetic cluster within the SARS-CoV-2 clade of sarbecoviruses, which might be designated the Japanese clade of bat sarbecoviruses (Figure 3B).

**Table 2.**
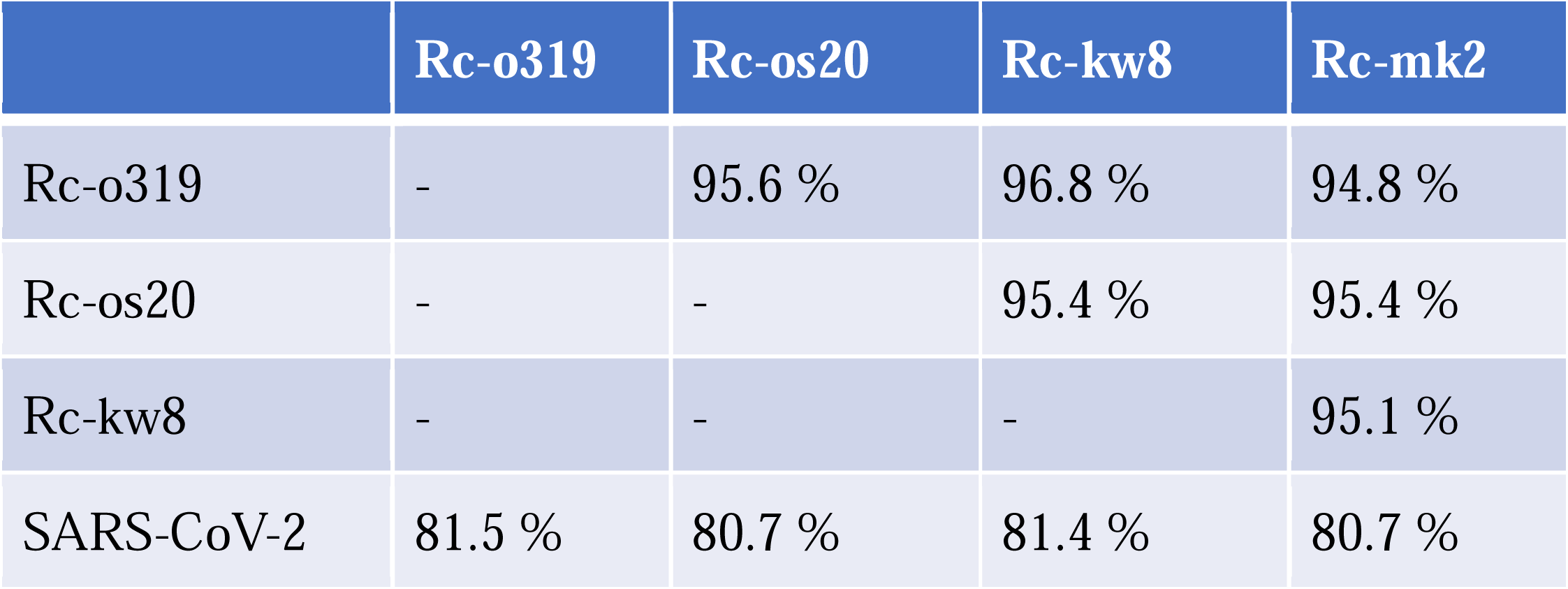
Full genome nucleotide identity among Japanese isolates

**Figure 3.**
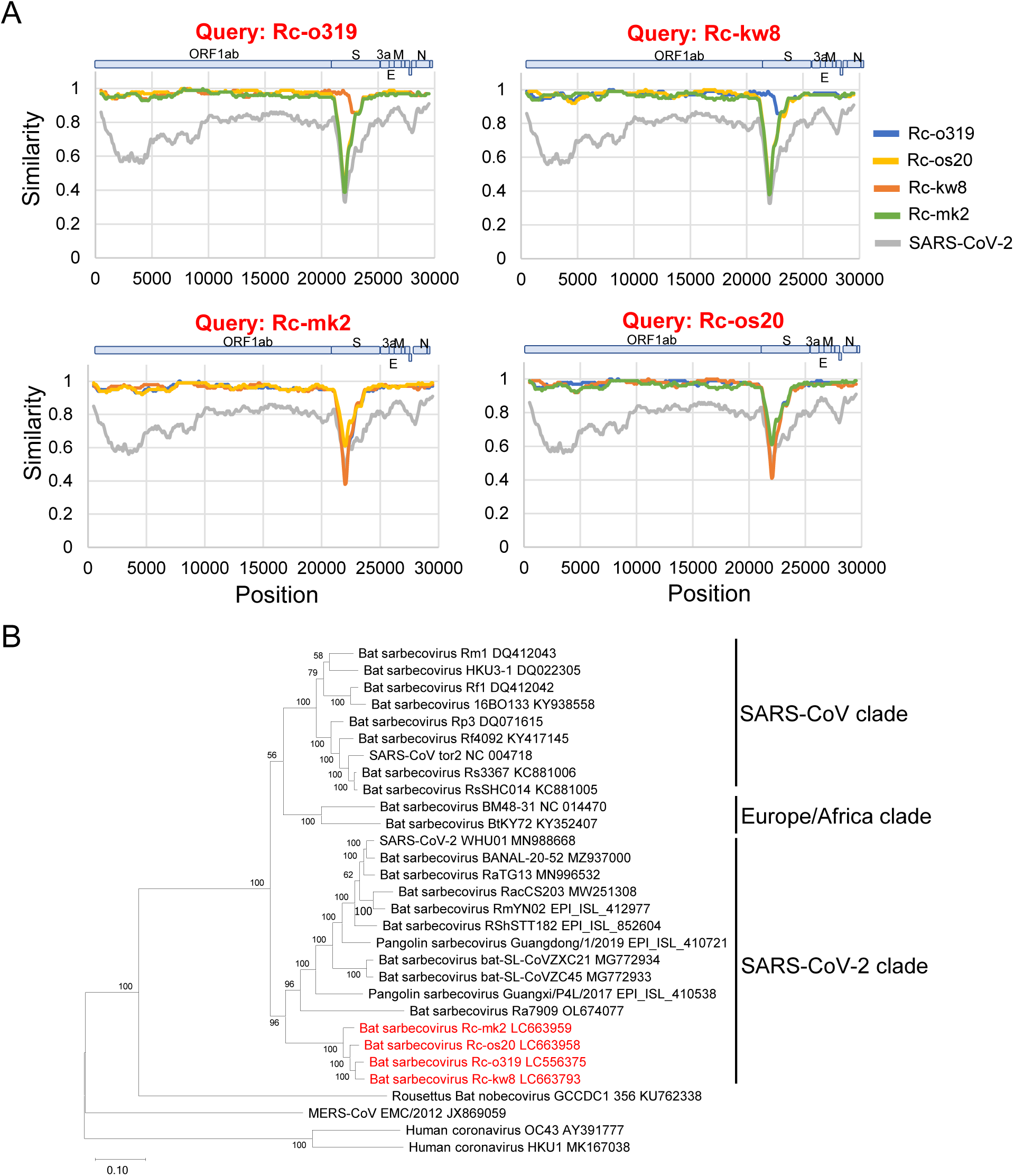
Genetic analysis of bat sarbecovirus isolates in Japan. (A) Similarity plot analysis of isolates was performed using the full-length genome sequence of Rc-o319, Rc-os20, Rc-kw8, or Rc-mk2 as query. SARS-CoV-2 virus was used as a reference. (B) Phylogenetic tree of bat sarbecoviruses was generated using the full genome nucleotide sequences with the maximum-likelihood analysis combined with 500 bootstrap replicates. Red text indicates the isolates in this study. Bootstrap values are shown above and to the left of the major nodes. Scale bars indicate nucleotide substitutions per site.

### Growth kinetics of Japanese isolates in cell culture

We aligned the receptor binding motif of the S protein of Japanese isolates with that of other sarbecoviruses (Figure 4A). We observed that all isolates had a 9-amino acid deletion in this motif, as previously observed in Rc-o319, and possessed relatively conserved residues with Rc-o319. Therefore, we assumed that the use of hACE2 as an entry receptor is unlikely in these new Japanese isolates. To test this hypothesis, we compared the replication of Japanese isolates and that of a control SARS-CoV-2 (B.1.1.7, alpha variant) in Vero-RcACE2, Vero-hACE2, Vero-ACE2KO, and Vero/TMPRSS2 cells. We found that the 4 bat isolates replicated well only in Vero-RcACE2, whereas did not replicate in Vero/TMPRSS2, Vero-hACE2, or Vero-ACE2KO cells, suggesting their RcACE2-dependent infectivity. In contrast, we noticed that SARS-CoV-2 replicated efficiently in Vero/TMPRSS2, Vero-RcACE2, and Vero-hACE2 cells, but not in Vero-ACE2KO cells (Figure 4B), suggesting multiple ACE2-dependent infectivity including that of *R. cornutus*. These data suggested that Japanese bat isolates use only RcACE2 as a receptor, showing narrow host specificity.

**Figure 4.**
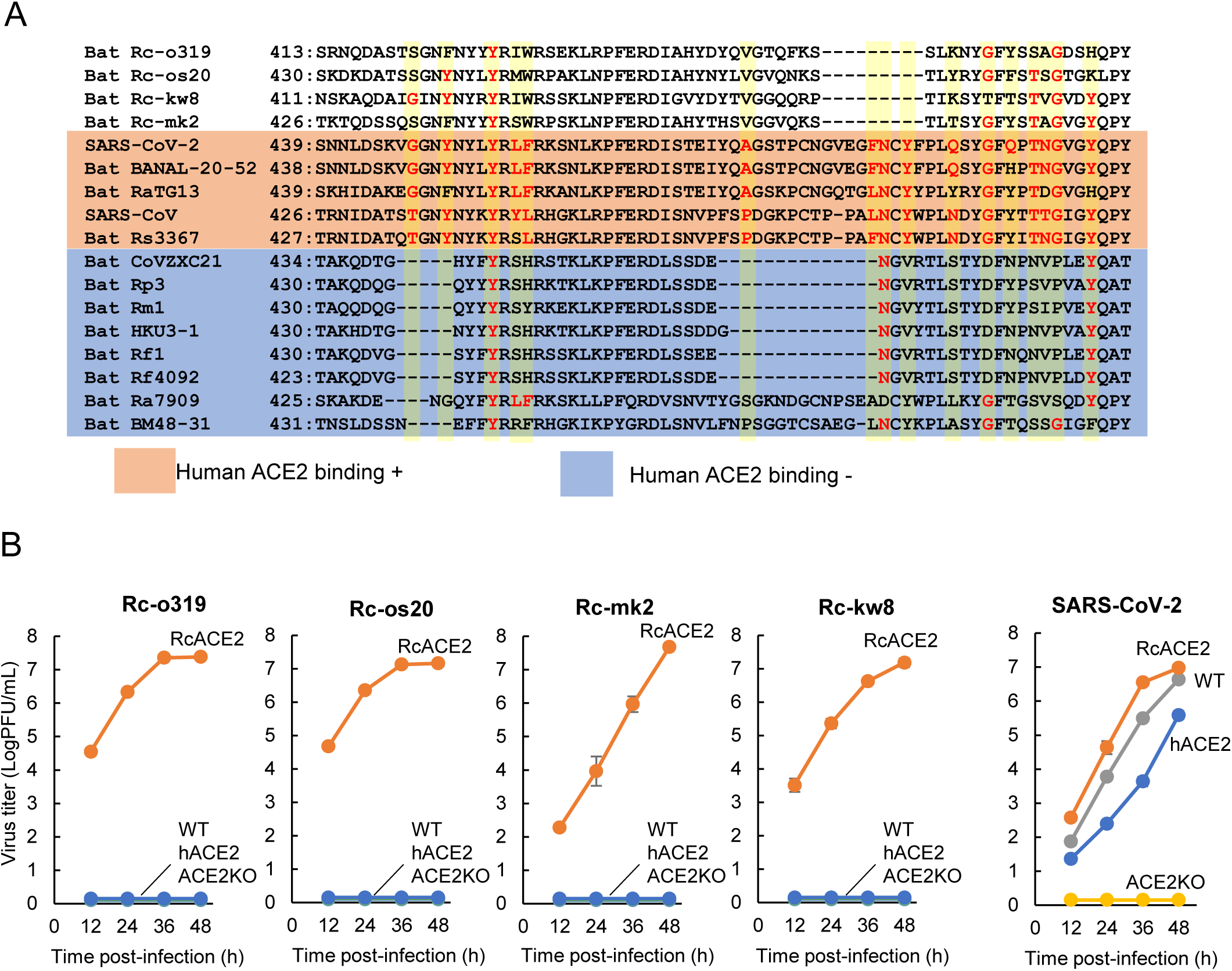
Receptor binding motif (RBM) and growth kinetics of bat sarbecovirus isolates. (A) Alignment of the RBM sequence of S proteins. Amino acid positions of RBMs contacting human ACE2 identified in SARS-CoV-2 are shaded in yellow, while residues identical to SARS-CoV or SARS-CoV-2 are shown in red. Strains that are capable to bind to human ACE2 are shaded in orange, whereas those that are incapable to bind to human ACE2 are given in blue. (B) Growth kinetics of isolates. Bat isolates Rc-o319, Rc-os20, Rc-mk2, and Rc-kw8 or SARS-CoV-2 (B.1.1.7) were inoculated into Vero/TMPRSS2 (WT), Vero-RcACE2 (RcACE2), Vero-hACE2 (hACE2), or Vero-ACE2KO (ACE2KO) cells at an MOI of 0.01. The culture supernatants were collected at the indicated time points, and viral titers were determined using a plaque assay. Data were reported as the mean titer with standard deviations from 3 independent experiments.

## Discussion

In this study, we successfully isolated 4 bat sarbecoviruses from *R. cornutus* bats in distinct locations in Japan, using Vero-RcACE2 cells. The origin of SARS-CoV-2 is thought to be a sarbecovirus from *Rhinolophus* spp. bats in southern China and the Indochinese peninsula, as bat viruses detected in these regions have been found to have high homology with SARS-CoV-2 and to be able to efficiently bind to human ACE2 (1, 3, 7, 8, 13-15). In contrast, the Japanese bat sarbecovirus isolates were phylogenetically distant from SARS-CoV-2, compared with bat sarbecoviruses from southern China and the Indochinese peninsula, despite belonging to the SARS-CoV-2 clade. In addition, the Japanese bat isolates were unable to use hACE2 as a cell entry receptor, thus making it unlikely to be one of the ancestors of SARS-CoV-2. However, phylogenetic analysis indicated that the Japanese viruses were positioned near a branching node between the SARS-CoV and SARS-CoV-2 clades, suggesting that they might be related to the common ancestor of SARS-CoV and SARS-CoV-2. These findings suggested that characterization of bat viruses might provide an understanding of the mechanism and the potential of bat sarbecoviruses to overcome host barriers.

Because of the pilot study aspect of this study, the number of samples and the bat species studied were limited. Four species of *Rhinolophus* bats inhabit Japan: *R. cornutus, R. ferrumequinum, R. pumilus*, and *R. perditus*. The former 2 species are widely distributed, exhibiting a major population in Japan, whereas the latter 2 species are mainly confined in Okinawa prefecture, which is located far southwest of the main island of Japan. In this study, the bat sarbecoviruses were isolated from *R. cornutus* but not from *R. ferrumequinum*. Since the virus has been detected from *R. ferrumequinum* in China (5), it is expected that increasing the number of samples will clarify whether *R. ferrumequinum*-associated virus existents in Japan. Although there have been no reports of sarbecovirus detection in *R. pumilus* and *R. perditus*, it is likely that both species might harbor sarbecoviruses with different genetic characteristics from the isolates in our study because of their different niche. Further epidemiological surveys are needed to confirm this hypothesis.

Virus isolates can be used to elucidate their biological characteristics, including pathogenicity and antigenicity. To this date, all detected bat sarbecoviruses are classified into 2 groups based on receptor selectivity: hACE2-binding and non-hACE2-binding types. In particular, hACE2-binding type bat sarbecoviruses were isolated from African green monkey Vero cells, whose ACE2 molecule might be functionally related to hACE2 (9). In contrast, to the best of our knowledge, non-hACE2-binding type bat sarbecoviruses have not been isolated. Comparative analysis between hACE2-binding and non-hACE2-binding viruses would facilitate an understanding of the factors that determine receptor specificity of bat sarbecoviruses.

Both the NTD and RBD of the S protein were highly variable among sarbecoviruses (25-28), probably due to escape from immune pressure, as both regions contain viral neutralizing epitopes (29-31). *Rhinolophus* spp. bats are relatively short-distance migrants (32, 33); hence, the migration of individuals between bat groups is infrequent. Most genome sequences, except for the RBD- and NTD-coding regions of the S gene, were highly conserved among Japanese strains, suggesting that Japanese sarbecoviruses diverged relatively recently from the undefined ancestral virus, and consecutively rapidly accumulated mutations in the NTD and RBD regions due to strong selection pressure. Several studies have been reported that in addition to sarbecoviruses *Rhinolophus* bats harbor alphacoronaviruses (8, 34), suggesting that a virus could undergo major changes by recombination with other coronaviruses in the NTD/RBD regions, leading to the emergence of a novel coronavirus with zoonotic potential.

Various sarbecoviruses lacking ORF8 protein have been identified in bats (6, 35) and humans (36, 37). Although the function of ORF8 protein in SARS-CoV-2 is not fully understood, it might act as an inhibitor of IFN-I signaling (38) and a downregulator of MHC-I (39). Hence, an ORF8-deleted virus shows lower pathogenicity in humans (40). Among our bat isolates, 2 strains possessed the ORF8 gene, but their homology to that of SARS-CoV-2 was very low (approximately 27.5 % in amino acid sequence), suggesting functional differences. Likewise, no appreciable difference was detected in growth in cell culture between viruses possessing or lacking ORF8 protein (e.g., Rc-o319 versus Rc-os20). Further studies are required to clarify the biological characteristics of ORF8 protein in bat sarbecoviruses.

In conclusion, we isolated bat sarbecoviruses from *R. cornutus* in several locations in Japan that were phylogenetically positioned in the same cluster of the SARS-CoV-2 clade. These isolates did not replicate in hACE2-expressing cells, suggesting low potential for human infection. However, sarbecoviruses might mutate and infect humans via intermediate hosts in wildlife or livestock. Therefore, epidemiological studies of sarbecoviruses in wildlife, including bats, need to be conducted on a long-term basis for risk assessment of their zoonotic potential.

Dr. Murakami is an associate professor at the Graduate School of Agricultural and Life Sciences, University of Tokyo, Tokyo, Japan. His research interests include epidemiologic and molecular biological studies of animal viruses, including coronaviruses and influenza viruses.

## Acknowledgements

This work was supported by the Japan Agency for Medical Research and Development (AMED) under the grant number JP21fk0108615. We thank Ms. Satomi Kato for the technical assistance.

